# Loss of Protein Kinase D2 Activity Protects Against Bleomycin-induced Dermal Fibrosis in Mice

**DOI:** 10.1101/2021.10.02.462861

**Authors:** Liping Chen, Jinjun Zhao, Yapeng Chao, Adhiraj Roy, Wenjing Guo, Jiabi Qian, Wanfu Xu, Zhe Xing, Xiaoju Lai, Binfeng Lu, Fan Deng, Q. Jane Wang

**Author notes:** These authors are corresponding authors **Address correspondence to:** Q. Jane Wang, Department of Pharmacology and Chemical Biology, University of Pittsburgh, Fan Deng, Ph.D., Department of Cell Biology, School of Basic Medical Sciences, Southern Medical University. These authors contributed equally to this work (Liping Chen, Jinjun Zhao).

## Abstract

**Background:** Dermal fibrosis occurs in many human diseases, particularly systemic sclerosis (SSc) where persistent inflammation leads to collagen deposition and fiber formation in skin and multiple organs. The family of protein kinase D (PKD) has been linked to inflammatory responses in various pathological conditions, however, its role in inflammation-induced dermal fibrosis has not been well defined. Here, using a murine fibrosis model that gives rise to dermal lesions similar to those in SSc, we investigated the role of PKD in dermal fibrosis in mice lacking PKD2 activity.

**Methods:** Homozygous kinase-dead PKD2 knock-in mice (PKD2^SSAA/SSAA^-KI) were obtained through intercrossing mice heterozygous for PKD2^S707A/S711A^ (PKD2^SSAA^). The wild-type and KI mice were subjected to repeated subcutaneous injection of bleomycin (BLM) to induce dermal inflammation and fibrosis. As controls, mice were injected with PBS. At the end of the experiment, mouse skin at the injection site was dissected, stained, and analyzed for morphological changes and expression of inflammatory and fibrotic markers. PKD-regulated signaling pathways were examined by real-time RT-qPCR and Western blotting. In a separate experiment, BLM-treated mice were administered with or without a PKD inhibitor, CRT0066101 (CRT). The effects of CRT on dermal fibrosis were analyzed similarly. The identity of the PKD expressing cells were probed using myeloid lineage markers CD45, CD68 in BLM-treated mouse tissues.

**Results:** Dermal thickness and collagen fibers of kinase-dead PKD2-KI mice were significantly reduced in response to BLM treatment as compared to the wild-type mice. These mice also exhibited reduced α-smooth muscle actin (α-SMA) and collagen expression. At molecular levels, both transforming growth factor β1 (TGF-β1) and interleukin-6 (IL-6) mRNAs were decreased in the KI mice treated with BLM as compared to those in the wild-type mice. Similarly, CRT significantly blocked BLM-induced dermal fibrosis and inhibited the expression of α-SMA, collagen, and IL-6 expression. Further analysis indicated that PKD2 was mainly expressed in CD45^+^/CD68^+^ myeloid cells that could be recruited to the lesional sites to promote the fibrotic process of the skin in response to BLM.

**Conclusions:** Knock-in of the kinase-dead PKD2 or inhibition of PKD activity in mice protected against BLM-induced dermal fibrosis by reducing dermis thickness and expression of fibrotic biomarkers including α-SMA, collagen, and inflammatory/fibrotic mediators including TGF-β1 and IL-6. PKD2 does this potentially through modulating the recruitment and function of myeloid cells in skin of BLM-treated mice. Overall, our study demonstrated a potential critical role of PKD catalytic activity in inflammation-induced dermal fibrosis.

## Introduction

Dermal fibrosis assoicates with a number of diseases, most notably Systemic Sclerosis (SSc) which is a complex multisystem connective tissue disease with high mortality (1). It has an incidence of nearly 20 patients per million per year (2, 3) with more than half of patients eventually dying as a direct consequence of the disease (4, 5). Based on the extent and distribution of skin involvement, SSc can be classified into two main subtypes: limited cutaneous and diffuse cutaneous SSc. The primary pathological features of SSc are aberrant immune activation, vasculopathy, and tissue fibrosis with considerable clinical heterogeneity. Following the early inflammatory stage of SSc, tissue deposition of collagen-rich scar occurs, disrupting the normal architecture of the skin and leading to dysfunction and eventual failure of the skin, lung, and other organs (6). The treatments for SSc are often symptomatic and have limited effectiveness.

Inflammation and fibrosis are intimately coupled during the progression of tissue fibrosis. Inflammatory responses are not only triggered during early stage of tissue injury, but also accompany the development of severe fibrosis at later stages. Advances in the past few years have led to the identification of many important inflammatory and fibrotic mediators, such as TGF-β1 and IL-6, which play key roles in the pathogenesis of dermal fibrosis. Several experimental dermal fibrosis models have been developed that bore resemblance that occurring in human SSc (7). Among them, BLM-induced fibrosis model is the most widely used in SSc studies. It is well established that mice receiving daily subcutaneous injection of BLM for 4–6 weeks develop dermal fibrosis with autoantibody production, and immune infiltration, which mimicked the biochemical and histologic features of human SSc (8, 9). This model was used in this study to investigate the role of protein kinase D2 (PKD2) in dermal fibrosis.

PKD belongs to the Ca^2+^/calmodulin-dependent protein kinase super-family, with three different isoforms being identified: PKD1, PKD2 and PKD3 (10) with PKD1 being the most extensively studied isoform. PKD plays an important role in a variety of cellular processes including cell proliferation, migration, secretion, and inflammation (11). Its involvement in the inflammatory process has been implicated in several studies. PKD is required for the production of proinflammatory cytokines and chemokines in tumor cells (12), immune cells (13-15) and endothelial cells in response to various stimuli (16). PKD has been implicated in the following inflammatory conditions: pancreatitis, hypersensitivity pneumonitis (13), autoimmune diseases, allergic inflammatory diseases, viral infection, airway inflammation, and Sjogren’s syndrome-related inflammation (11, 13, 15, 17-23). PKD has also been implicated in heart and lung fibrosis (24, 25). PKD is increased and activated in lung epithelial cells and macrophages in idiopathic pulmonary fibrosis and may participate in the pathogenesis of this disease (25). However, little is known about the precise role of PKD in dermal fibrosis.

In this study, knock-in of kinase-dead PKD2 or inhibition of PKD activity prevented dermal fibrosis in a BLM-induced fibrosis mouse model. We showed that the KI mice or PKD inhibitor-treated mice exhibited reduced dermal thickness, decreased α-SMA expression, collagen deposition, and production of pro-inflammatory/pro-fibrotic mediators in response to BLM treatment *in vivo*. We further identified PKD-expressing cells in dermis of BLM-treated mouse skin. Our study demonstrated a potential critical role of this kinase family in BLM-induced dermal fibrosis through modulating immune infiltration and cytokine production.

## Materials and Methods

### Reagents

These antibodies are obtained from Cell Signaling Technology (Danvers, MA): PKD1, PKD2, PKD3 (D57E6 Rabbit mAb), p-PKD1(Ser916), CD45, CD68, and α-Smooth Muscle Actin. The p-PKD2 (Ser876) and eBioscience™ 1X RBC Lysis Buffer were obtained from Thermo Fisher (Pittsburgh, PA). TrueBlack Lipofuscin Autofluorescence Quencher was from Biotium. Donkey anti-Rabbit IgG (H+L) Highly Cross-Adsorbed Secondary Antibody, Alexa Fluor 488, and Donkey anti-Rabbit IgG (H+L) Highly Cross-Adsorbed Secondary Antibody, Alexa Fluor 568, and Mouse M-CSF Recombinant Protein were from Invitrogen (Carlsbad, CA). Lipopolysaccharides from Escherichia coli O111:B4 was purchased from Sigma (St Louis, Mo). Bleomycin hydrochloride was from Nippon Kayaku (Tokyo, Japan).

### Breeding of PKD2-Knock-In Mice

Catalytic deficient heterozygous PKD2^S707A/S711A/WT^ -knock-in (PKD2^SSAA/WT^-KI) mice on a C57BL/6 background were obtained from Jackson Laboratory (Bar Harbor, ME). Male PKD2 wild-type mice and PKD2^SSAA/WT^-KI mice of 6 wks old (body weigh ∼20 g) were maintained with food and water ad libitum and housed in specific pathogen-free condition. Mouse genotyping was carried out by PCR amplification of genomic DNA (26). Homozygous PKD2^SSAA/SSAA^-KI mice were obtained by intercrossing heterozygous PKD2^WT/SSAA^ mice (27).

### Mouse model of bleomycin-induced dermal fibrosis

Healthy wild-type and PKD2^SSAA/SSAA^-KI mice were selected to match in gender, age (6-8 wks) and weight. The mice were divided into four groups: two groups of PKD2^WT/WT^ mice receiving vehicle (1xPBS) or BLM: PKD2^WT/WT^ -PBS and PKD2^WT/WT^-BLM; two groups of mutant mice receiving PBS or BLM: PKD2^SSAA/SSAA^-KI-PBS and PKD2^SSAA/SSAA^-KI-BLM. Bleomycin was prepared by dissolving in 1xPBS at a concentration of 1 mg/mL, followed by sterilization by filtration (28). To induce dermal fibrosis, BLM (100 µL at 1 mg/mL) was injected subcutaneously to an shaved area (∼3 cm^2^) on the back of the mice daily for 4 wks as described by *Yamamoto et al*. (9) and *Shun Kitaba et al*. (29), while the control mice received 100 µL of PBS instead. This model is characterized by infiltration of mononuclear cells into the lesional skin and thickening of the dermis that is maintained for at least 6 wks after cessation of treatment (8). To measure lymphocyte infiltration, wild-type and PKD2 knock-in mice were injected subcutaneously with BLM (100 µL at 1 mg/mL) for 1 week. The skin tissues at the lesion site were collected for IHC analysis.

For PKD inhibitor treatment, healthy female wild-type C57BL/6 mice were divided into three groups receiving vehicle (1xPBS), BLM, and BLM plus CRT0066101 (CRT). CRT was administered orally at 80 mg/kg in a 5% dextrose saline solution every day for 4 weeks. Mouse weight was measured once per week.

### Mouse tissue sample preparation

To collect tissues, mice were anesthetized after having completed bleomycin treatment for 7 or 28 days. After removing hair at the bleomycin injection site, the dorsal skin was carefully dissected by RNA-free surgical scissors and divided into three parts, one was fixed in neutral buffered formalin (10% NBF) for paraffin-embedding, sectioning, hematoxylin and eosin (H&E) staining, and immunohistochemistry (IHC) analysis, one was embedded in OCT and flash frozen for sectioning and immunofluorescence (IF) staining, and the remaining, after eliminating the subcutaneous fat layer, was snap-frozen in liquid N2 and stored in -80°C for Western blotting and RT-QPCR analysis.

### Western blot analysis

The skin tissue samples were ground in liquid nitrogen and lysed in lysis buffer. Lysates were separated by SDS-PAGE and transferred onto nitrocellulose membrane. The membranes were pre-blotted with 5% non-fat dry skim milk plus 2% bovine serum albumin (BSA) in PBS with 0.05% Tween 20 (PBST) for 1 h and then incubated with anti-Collagen I-α2 (Proteintech, 1:1000), anti-α-SMA (Proteintech, 1:1500), anti-PKD1 (Santa Cruz, 1:1000), anti-p-PKD1 (ser744/748) (Cell signaling technology, 1:1000), anti-PKD2 (ABclonal, 1:1500), anti-p-PKD2 (ser876) (Sigma, 1:1000), anti-PKD3 (Cell signaling technology, 1:1500), anti-STAT3 (Cell signaling technology, 1:2000), anti-p-STAT3 (Tyr705) (ABclonal, 1:1500), anti-α-tubulin (Beijing Ray Antibody Biotech, China, 1:4000) antibodies overnight at 4°C, washed with PBST and followed by secondary antibodies. Finally, proteins were detected by the enhanced chemiluminescence method (SuperSignal West Pico Chemiluminescent Substrate, Pierce, Rockford, IL).

### RNA extraction and real time quantitative RT-qPCR analysis

Total RNA was extracted from tissues using TRIzol reagent (Invitrogen) following the manufacture’s protocol. Briefly, reverse transcription and real time RT-qPCR analysis were carried out using the ALL-in-ONE First-Strand cDNA Synthesis Kit and All-in-One qPCR Mix (GeneCopoeia) and the SsoFast EvaGreen Supermix according to the manufacturer’s protocols on CFX96 Real-Time PCR Detection System (Bio-Rad, Richmond, CA). Data were normalized using GAPDH as an internal control. All primers for real time RT-qPCR were purchased from Invitrogen Life Technologies and sequences of the primers are listed in Supplementary Table 1.

### Bone Marrow-Derived Macrophages (BMM) Preparation

Femur and tibia of wild-type C57BL/6 mouse were obtained from ice. Bone marrow was flushed with 10mL RPMI-1640 media and centrifuged at 1200 rpm for 5 min. The supernatant was discarded and cell pellets were treated with eBioscience™ 1× RBC Lysis Buffer. After washing, the cells were cultured in 10% FBS of RPMI-1640 supplemented with mouse M-CSF (Thermo Fisher). Medium was replenished every 3 days and day-6 macrophages were used for experiment.

### Immunofluorescence Staining

Slides containing frozen tissue sections were washed three times with 1X PBS, fixed in 4% paraformaldehyde at room temperature for 30 minutes. After washing, the sections were incubated in 5% normal goat serum containing 0.3% Triton X-100 for 1 hr at room temperature, then probed with the primary antibodies diluted in antibody dilution buffer (1% BSA, 0.3% Triton X-100 in 1X PBS) overnight at 4 °C. Thereafter, after washing, the sections were incubated with secondary antibodies diluted in antibody dilution buffer for 1 hr at room temperature, followed by counterstaining with DAPI (1 μg/ml). The coverslips were mounted in ProLong Gold antifade reagent (Invitrogen), and analyzed using an Olympus Fluoview (FV1000) confocal microscope using 60 X/1.45 objectives. Prior to mounting, the tissue sections were treated with 1X True Black Lipofuscin for 1 min at room temperature to decrease autofluorescence. The primary antibodies used were α-PKD2 (CST 8188), α-phospho PKD2Ser876 (Thermo PA5-64538). The secondary antibodies used were Alexa Fluor 488 donkey anti-rabbit IgG, Alexa Fluor 568 donkey anti-rabbit IgG, and Alexa Fluor 568 goat anti-mouse IgG. Macrophages were grown on poly-D-lysine coated coverslips and stained similarly as described above.

### Statistical analysis

Statistical analysis was performed by SPSS 21.0 software (SPSS, Chicago, IL) using independent sample t-test or one way-analysis of variance (ANOVA). Data are presented as mean ± S.D. A *p* value <0.05 was considered as significant.

## Results

### Knock-in of kinase-dead PKD2 blocked BLM-induced mouse dermal fibrosis

Homozygous PKD2^SSAA/SSAA^-KI mice were generated by replacing wild-type PKD2 alleles with mutant alleles that encode alanine substitutions of Ser707 and Ser711 (S707A/S711A) in the activation loop of PKD2 (Supplemental Fig. S1A). The phosphorylation of the two conserved serine residues in the activation loop is essential for PKD activity (30). This mouse model was first reported by Matthews et al. (26). The homozygous mice are viable, fertile, normal in size, and do not display any gross physical or behavioral abnormalities. Mouse genotyping confirmed the presence of mutated alleles in the KI mice (supplemental Fig. S1B) and Western blotting demonstrated absence of PKD2 activity measured by p-PKD2(ser876) antibody in various tissues (supplemental Fig. 1C).

Subcutaneous injection of BLM at 1 mg/mL daily for 28 days leads to dermal fibrosis in mice, a key feature of SSc (29). To investigate the role of PKD2 in this process, mice were randomized into four groups that received either vehicle (1xPBS at 100 uL/mouse, once daily) or BLM (BLM at 1 mg/mL, once daily) (n=9 mice): PKD2^WT/WT^-PBS, PKD2^WT/WT^-BLM, PKD2^SSAA/SSAA^-PBS and PKD2^SSAA/SSAA^-BLM (Fig. 1A). Dermal thickness is a crucial indicator of dermal fibrosis (31). Histopathological analysis of H&E stained skin sections from the four groups of mice showed that dermal thickness was increased over 2-fold in response to BLM in PKD2^WT/WT^ mice as compared to vehicle-treated mice, while this response was significantly reduced in the PKD2^SSAA/SSAA^-KI mice receiving BLM injection (Fig. 1C, 1B). Additionally, the effect was only observed in the dermal layer, but not in the epidermal layer, which agrees with other reports (32) (Fig. 1C-D). In summary, we showed that inactivation of PKD2 as in PKD2^SSAA/SSAA^-KI mice blocked dermal thickening in the model of BLM-induced dermal fibrosis in mice.

**Figure 1.**
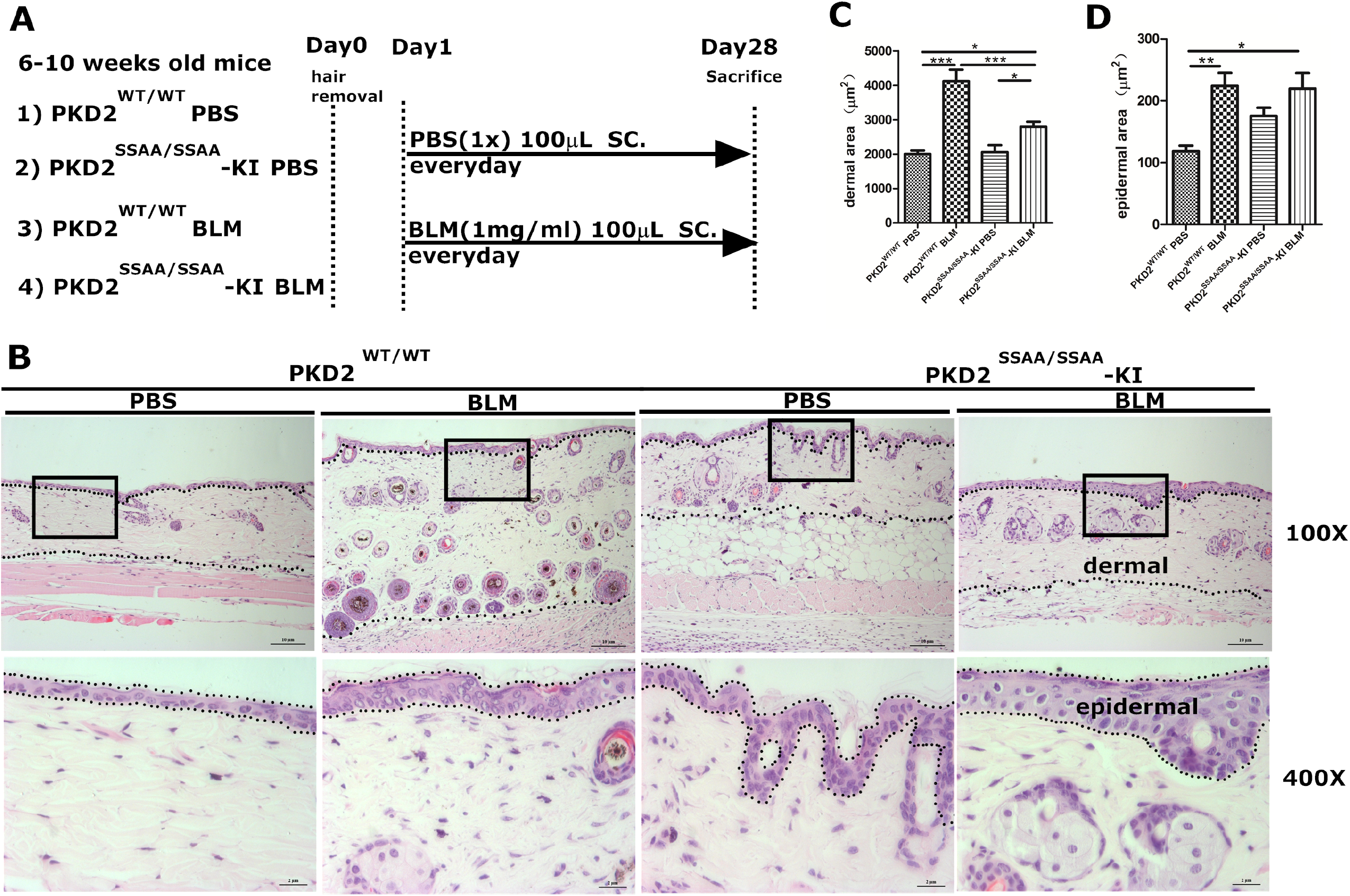
PKD2^SSAA/SSAA^-KI mice were resistant to BLM-induced dermal fibrosis. (**A**) A diagram depicting mouse model of BLM-induced fibrosis. Briefly, four groups of mice were subjected to daily subcutaneous injection of 100µL PBS or BLM (1 mg/mL) for 28 days (n=9). (**B**) Representative images of H&E stained of skin sections from PKD2^WT/WT^ mice or PKD2^SSAA/SSAA^-KI mice injected with PBS or BLM. Regions between the dotted lines were the measured areas of the dermal layer or the epidermal layer. The black box indicated the position of 400x images. Original magnifications: ×100, scale display 10µm; ×400, scale display 2µm. (**C-D**) The dermal and epidermal thickness were measured by Image J software. Data are shown as the means±SD of triplicate determinations per 100x image from three mice per group. Statistical significance was analyzed by one-way ANOVA with multiple comparisons. *p<0.05, ***p<0.001.

### Kinase-dead PKD2 significantly abated collagen fiber expression in bleomycin-treated mouse dermis

Fibrosis is characterized by excessive deposition of extracellular matrix components such as collagens and fibronectin (33, 34). We examined collagen fibers by Masson-Trichrome staining. As shown in Fig. 2, the dermal layers in PKD2^SSAA/SSAA^-BLM group were arranged normally, however, the collagen fiber staining was weaker in the PKD2^SSAA/SSAA^ mice as compared to those in the PKD2^WT/WT^ mice treated with BLM (Fig. 2A). Quantitative analysis indicated that the BLM-induced collagen fiber deposition was greatly reduced in PKD2^SSAA/SSAA^-BLM group as compared to the control PKD2^WT/WT^-BLM group (Fig. 2B). Thus, PKD2 catalytic activity was required for BLM-induced collagen deposition in the dermis.

**Figure 2.**
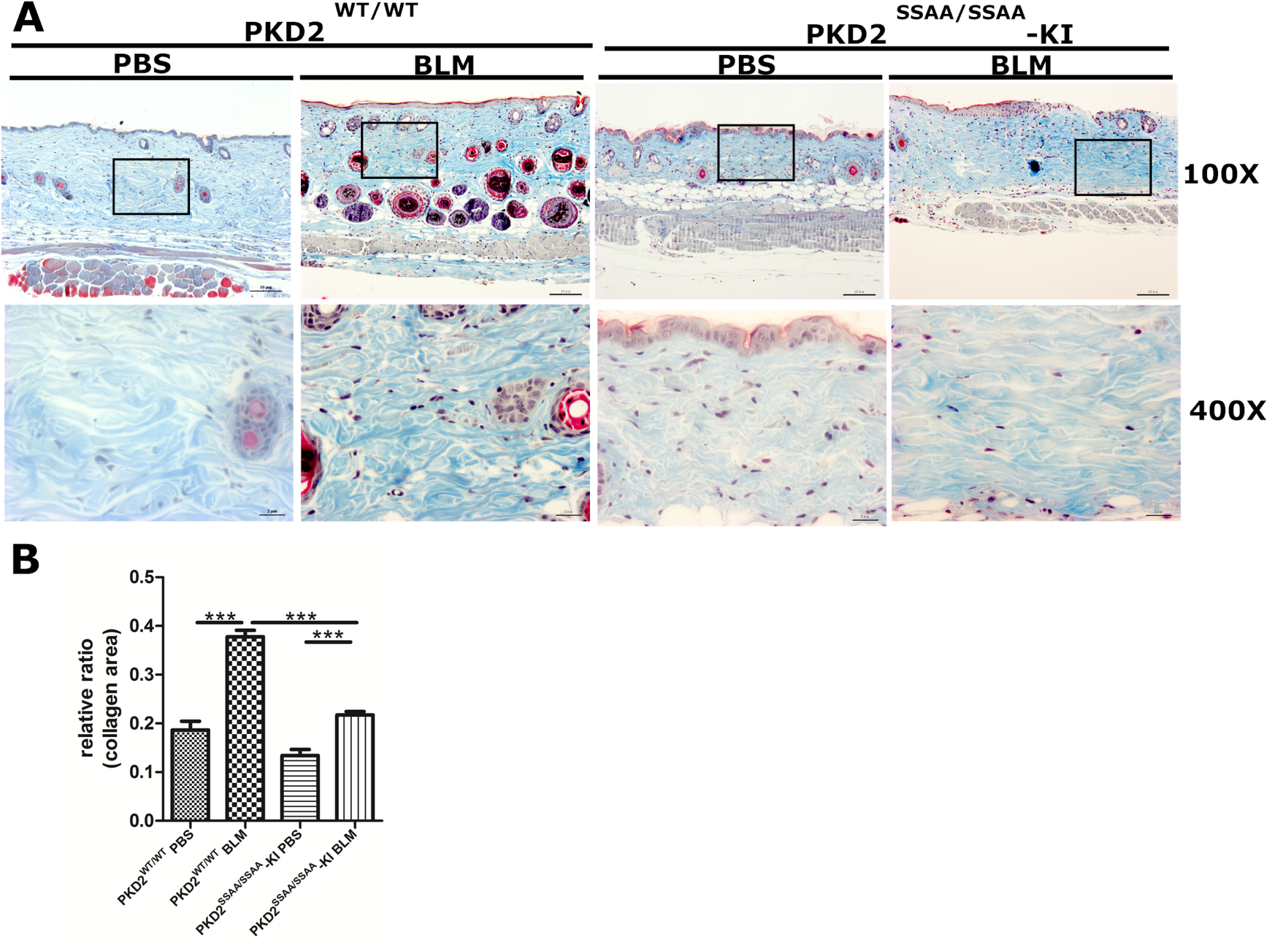
PKD2^SSAA/SSAA^-KI mice showed reduced collagen fibers in dermis after BLM treatment. (**A**) Masson-Trichrome staining of skin sections from PKD2^WT/WT^ or PKD2^SSAA/SSAA^-KI mice injected with PBS or BLM. Collagen fibers were stained blue and the nuclei were stained black. The black box indicated the position of 400x images. Original magnifications: ×100, scale display 10µm; ×400, scale display 2µm. (**B**) Collagen fibers relative ratio (%) of four groups measured by Image J software and analyzed by one-way ANOVA with multiple comparisons. **p<0.01,***p<0.001.

### The levels of α-SMA, collagen, and pro-fibrotic/pro-inflammatory cytokines were significantly reduced in the dermis of bleomycin-treated KI mice

Myofibroblasts, a key player in fibrotic process, are differentiated forms of fibroblasts that have features of both smooth muscle cells and fibroblasts, and are characterized by the expression of α-smooth muscle actin (α-SMA) (35). Here, we examined the expression of α-SMA and collagen in the PKD2^WT/WT^ and PKD2^SSAA/SSAA^ mice. As shown in Fig. 3A, mRNA levels of α-SMA decreased in PKD2^SSAA/SSAA^ mice treated with or without BLM as compare to those in PKD2^WT/WT^ mice. Additionally, mRNA expressions of collagen I, collagen IV-α1 and collagen IV-α2 were also reduced in BLM- and vehicle-treated PKD2^SSAA/SSAA^ mice as compared to those in the PKD2^WT/WT^ mice (Fig. 3A). Importantly, the levels of two main proinflammatory and profibrotic cytokines, IL-6 and TGF-β1, induced by BLM in PKD2^WT/WT^ mice were also significantly reduced in PKD2^SSAA/SSAA^ mice treated with or without BLM (Fig. 3B). It should be noted that although loss of PKD2 activity did not abolish BLM-induced α-SMA, collagens, TGF-β1, and IL-6 expression, the fold inductions of these genes upon BLM treatment in PKD2^SSAA/SSAA^ mice were lower as compared to those in PKD2^WT/WT^ mice, especially in terms of TGF-β1, and IL-6 (Fig. 3B). Thus, PKD2 activity is required for the induction of proinflammatory and profibrotic cytokines in BLM-induced dermal fibrosis.

**Figure 3.**
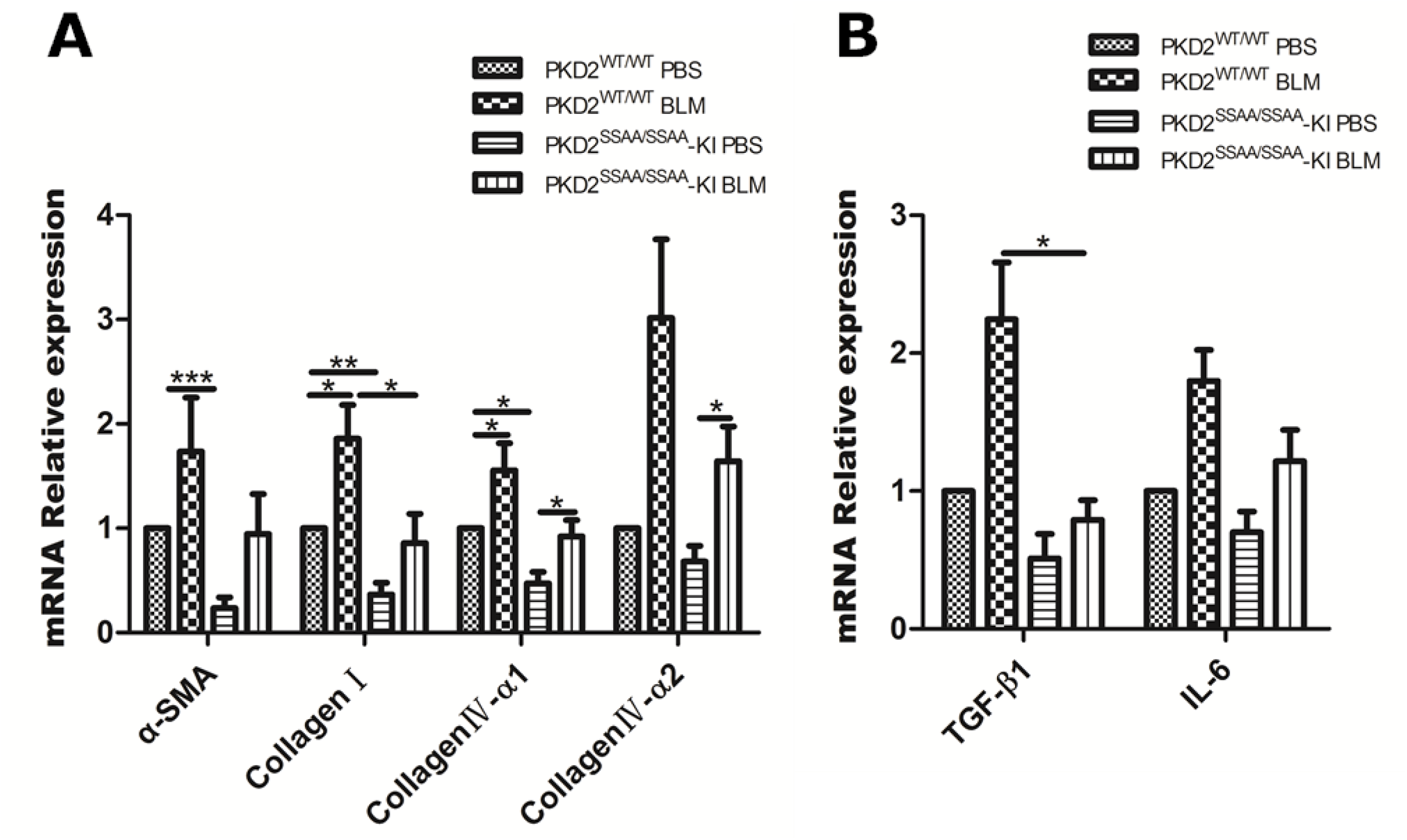
Analysis of α-SMA, collagen, TGF-β1 and IL-6 genes expression in skin of PKD2^WT/WT^ or PKD2^SSAA/SSAAA^-KI mice treated with BLM. (**A-B**) The total RNA of skin samples from indicated groups were extracted and mRNA levels of α-SMA, collagen I, collagen IV-α1, collagen IV-α2, TGF-β1 and IL-6 were measured by real time RT-qPCR with GAPDH as internal control. Data represent the means±SD of three mice, and were analyzed by one-way ANOVA with multiple comparisons. *p<0.05,**p<0.01,***p<0.001

Further analysis indicated that BLM failed to induce α-SMA protein expression in PKD2^SSAA/SSAA^-BLM mice (*p*<0.01) (Fig. 4A-B). Similarly, the protein level of collagen I-α2 was also decreased in PKD2^SSAA/SSAA^ mice treated with BLM as compared to the control group (Fig. 4A-B). In contrast, the levels of PKD1, 2, and 3 proteins and transcripts remained constant in all four groups of mice (Fig. 4A, 4C), while no p-S^876^-PKD2 was detected in dermis and epidermis of PKD2^SSAA/SSAA^ mice compared with those in the PKD2^WT/WT^ mice (Fig. 4A). Mouse skin sections were further analyzed for the presence of myofibrolasts, a key player in the fibrotic process after BLM treatment. As shown in Fig. 4D, there were significantly reduced number of α-SMA-positive cells in dermis of KI mice as compared to WT mice after 4 wks BLM treatment, which are in line with the Western blotting results (Fig. 4A). Taken together, our data indicate that PKD2 activity is necessary for α-SMA and collagen I-α2 expression in the dermis of BLM-treated mice.

**Figure 4.**
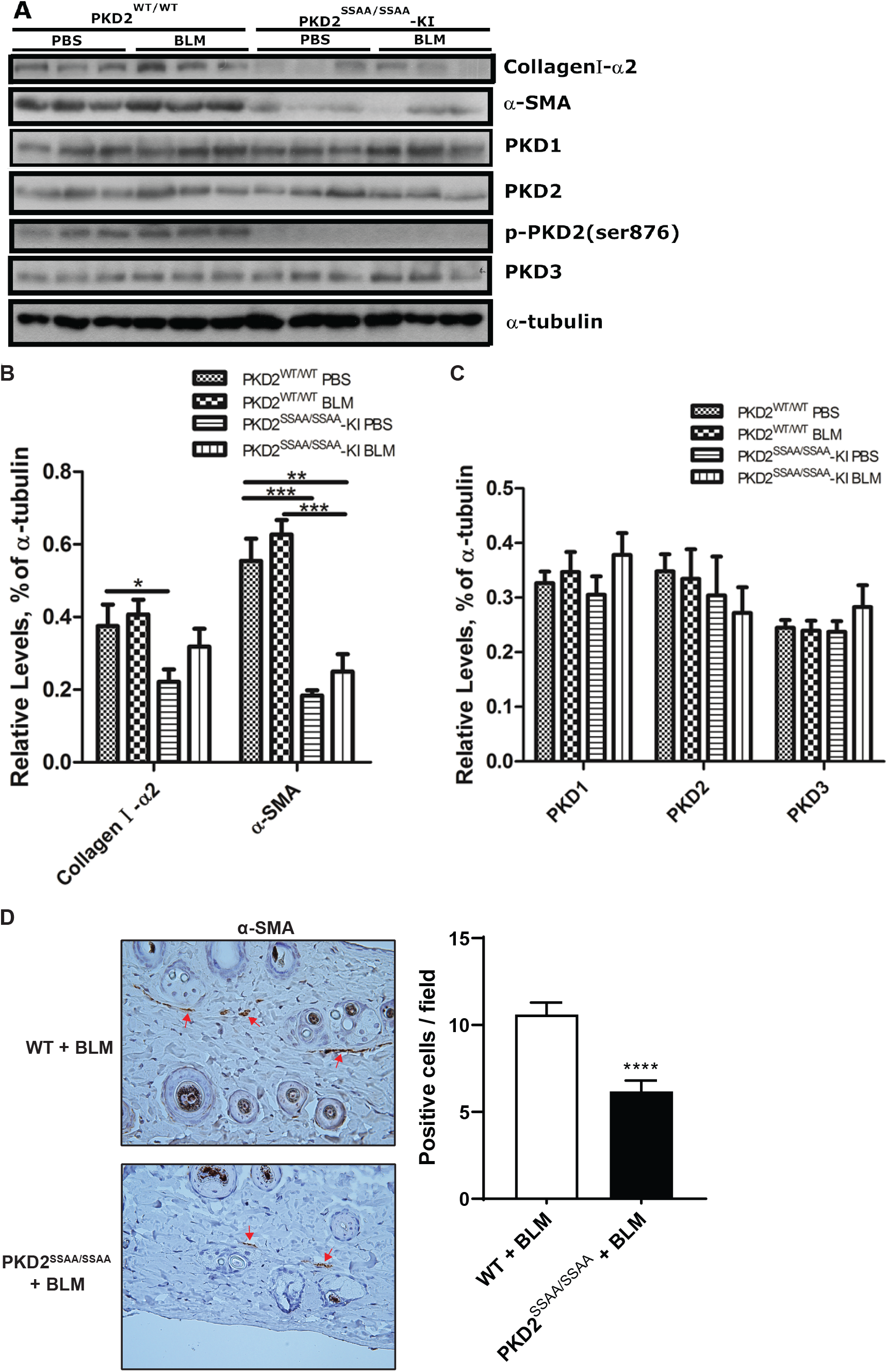
PKD2^SSAA/SSAA^-KI mice exhibited reduced collagen I-α2, α-SMA proteins and absence of PKD1 and PKD2 activities. (**A**) Western blotting analysis of the skin samples. KI and WT mice (n=9) were subjected to daily subcutaneous injection of 100µL PBS or BLM (1 mg/mL) for 28 days. Skin samples were lysed and analyzed by Western blotting for expression of α-SMA, collagenI-α2,p-PKD2 (S876), and nativePKD isoforms. α-tubulin was blotted as a loading control. (B)-(C) Corresponding densitometric quantification of collagen I-α2, α-SMA, PKD1, PKD2, and PKD3 levels in Western blots by Image J software and analyzed by one-way ANOVA with multiple comparisons. Data represent the means±SD from three separate experiments. *p<0.05, **p<0.01, ***p<0.001 (**D**) Reduced number of α-SMA-positive cells in mouse skin of BLM-treated KI mice as compared to WT mice. Mouse skin sections were analyzed by IHC using α-SMA antibody. *Right*, Representative IHC images, 40x; *Left*, Quantitative analysis of α-SMA-positive cells in the dermis from 6-10 random fields Mean±SD are shown. ****p<0.0001.

### Inhibition of PKD blocked BLM-induced dermal fibrosis in mice

To further corroborate our findings in mouse genetic models and exploit the therapeutic potential of targeting PKD, we investigated whether targeted inhibition of PKD using small molecule inhibitors has an impact on BLM-induced dermal fibrosis. CRT0066101 (CRT) is an orally available potent PKD inhibitor with demonstrated *in vivo* activity in multiple animal models (36-38). Female mice (7 wks old) were randomized into three groups (n=6) to receive daily subcutaneous injections of vehicle (1xPBS, 100 µL/mouse), BLM (1 mg/mL), and BLM + CRT (80 mg/kg, PO, once daily). H&E stained skin sections from the three groups of mice showed that while dermal thickness was significantly increased (∼1.5-fold) in response to BLM treatment as compared to vehicle-treated mice, CRT completely blocked this effect of BLM in the dermis (Fig. 5A-B). Additionally, as described above, the effect was only observed in the dermal layer, but not in the epidermal layer (Fig. 5B).

**Figure 5.**
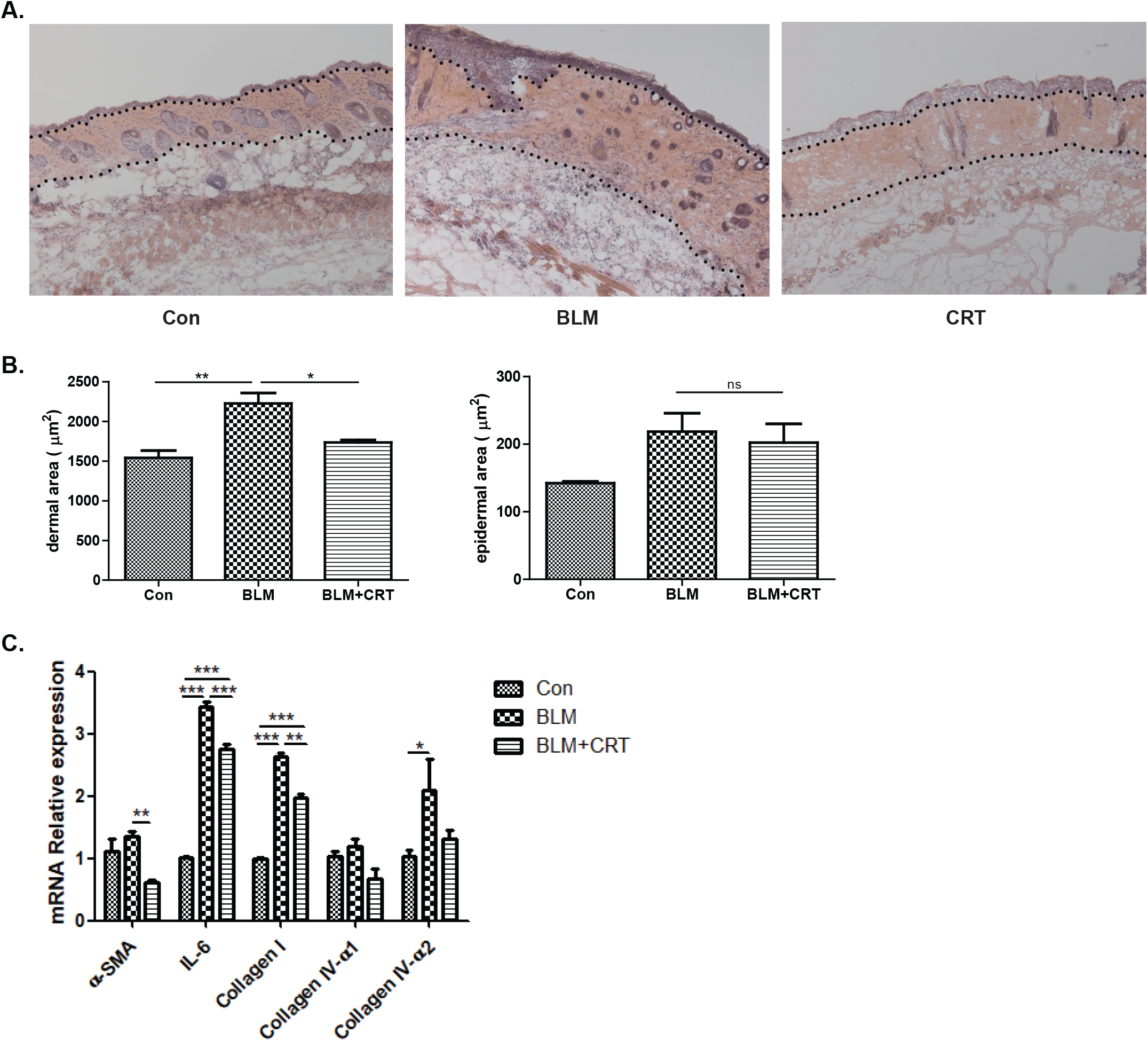
Inhibition of PKD blocked BLM-induced dermal fibrosis in mice. Healthy mice were randomized into three groups (n=6) for daily subcutaneous injection of PBS, BLM (1 mg/mL), and BLM + CRT (80 mg/kg) for 28 days. At the end of the treatment, skin around the injection site was harvested, sectioned, and subjected to H&E staining. (**A**) Representative images of H&E stained of skin sections from mice. Regions between the dotted lines were the measured areas of the dermal layer. Original magnification ×100. (**B**) The dermal and epidermal thickness were measured by Image J software. Data are shown as the means±SD of triplicate determinations per image from three mice per groups. Statistical significance was analyzed by one-way ANOVA with multiple comparisons. ns, no significant, *p<0.05, ***p<0.001. (**C**) Total RNA of skin samples from indicated groups were extracted and mRNA levels of α-SMA, collagen I, collagen IV-α1, collagen IV-α2, and IL-6 were measured by real time RT-qPCR with GAPDH as internal control. Data represent the means±SD of three mice, and were analyzed by one-way ANOVA with multiple comparisons. *p<0.05,**p<0.01,***p<0.001

The expression of α-SMA and collagens (collagen I, collagen IV-α1 and collagen IV-α2) was examined in the skin samples collected from the three groups of mice. As shown in Fig. 5C, mRNA levels of these genes were increased in response to BLM treatment, while significantly reduced by CRT, as compared to the vehicle-treated control group. Similarly, BLM-induced IL-6 was also significantly blocked by CRT (Fig.5C). Taken together, inhibition of PKD by CRT reversed BLM-induced dermal fibrosis in mice.

### Distinct patterns of PKD expression in mouse skin and the role of PKD2 in monocyte/macrophage infiltration in the lesional skin

To provide insights to how PKDs regulate skin function, we examined the distribution of the three PKD isoforms in skin of BLM-treated mice by IF staining. As shown in Fig. 6A, skin sections from mice treated with or without BLM showed distinction pattens of PKD1/2/3 distribution. PKD1 showed wide-spread distribution in both epidermis and dermis with predominant expression in the epidermis. After BLM treatment, the expression of PKD1 increased in the dermis. Interestingly, the adipose tissue layer in the hypodermis also showed prominent expression of PKD1 after BLM treatment. In contrast, PKD2 was found in discrete regions or sporadic cells/cell clusters within the dermis. In BLM-treated skin, there were increased PKD2-positive sporadic cells in the dermis. In comparison, PKD3 was mainly detected in hair follicle-like structures within the dermis before and after BLM treatment. Thus, PKD isoforms reside in distinct regions/cell types in the skin, suggesting they may play different roles in BLM-induced dermal fibrosis.

**Figure 6.**
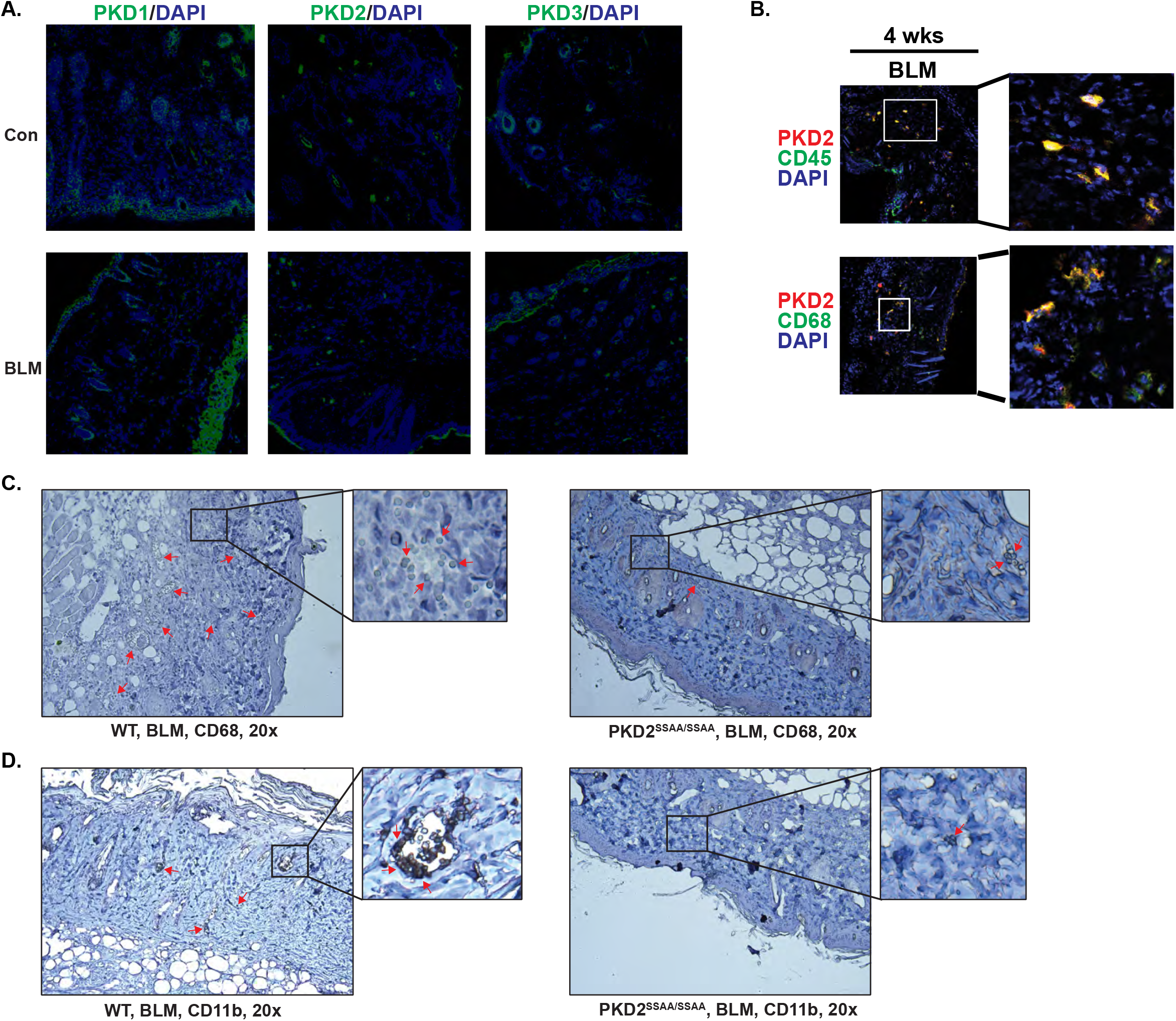
PKD2 resided in macrophages recruited to the lesional skin. (**A**) PKD1/2/3 distribution in mouse skin before and after BLM treatment. Frozen skin sections from mice treated with PBS (con) or BLM for 4 wks were subjected to IF staining using PKD1/2/3 antibodies. Representative merged images are shown. *green*, PKD1/2/3; *blue*, DAPI. Original magnification 100x. (**B**) PKD2 resided in CD45^+^/CD68^+^ myeloid cells in dermis of BLM-treated mice. Frozen tissue sections from 4 wks BLM-treated mice were co-stained with PKD2 or p-PKD2 (*red*) and CD45 or CD68 (*green*), and counterstained with DAPI (blue). Original magnification 100x. (**C-D**) Less CD68+ and CD11b+ macrophages were recruited to the dermis of BLM-treated KI mice as compared to WT mice. KI and WT mouse (n=3) were subjected to daily subcutaneous injection of 100µL BLM (1 mg/mL) for 7 days. Skin sections from 1 wks BLM-treated mice were analyzed by IHC using CD68 (C) and CD11b (D) antibodies and counterstained with hematoxylin. Original magnification x40. Representative images are shown.

To gain further understanding of the function of PKD2 in BLM-induced fibrosis, we sought to determine the identity of the PKD2 positive cells in skin of BLM-treated mice. Markers of myeloid lineage CD45 and CD68 were co-stained with PKD2 in skin sections from BLM-treated mice at 4 wks. As shown in Fig. 6B, PKD2-positive signals were nearly completely overlapped with those of CD45 and CD68. These data indicated that PKD positive cells likely belonged to the myeloid lineage. To determine if PKD2 activity affects immune cell infiltration, a critical event at the onset of inflammation leading to fibrosis, mouse skin tissue sections were obtained from KI and WT mice that have been treated with daily injection of BLM (1 mg/mL) for 7 days. Skin tissue sections were stained with CD68 and CD11b antibodies to reveal the presence of monocytes/macrophages, the main players in the inflammatory process. Our data demonstrated that the number of CD68+ and CD11b+ macrophages recruited to the dermis of BLM-treated KI mice was reduced as compared to that in WT mice, indicating that PKD2 activity is required for the recruitment of macrophages to the lesional site.

### PKD activity was required for the migration and cytokine production of bone merrow-derived macrophages

Our data implied that PKD2 was predominantly expressed in cells of the myeloid lineage which can reside or be recruited to the lesional skin in BLM-treatment mice. A major population of the myeloid cells marked by CD45/CD68 is macrophages, which have been implicated in the pathogenesis of dermal fibrosis in SSc. To determine if PKD affects the function of macrophages, we obtained bone merrow-derived macrophages (BMMs) from wild-type mice. The role of PKD in BMM migration and cytokine production, two critical functions of macrophages that contribute to the pathogenesis of dermal inflammation and fibrosis, were examined. As shown in Fig. 7A, BMM expressed all three PKD isoforms with PKD1 being most predominant. By co-staining for PKD1 and CD68, we confirmed that the BMM we obtained expressed the macrophage marker CD68, and PKD1 was well co-localized with CD68 (Fig. 7B), as observed in tissues sections. Next, BBM was stimulated with lipopolysaccharide (LPS), a potent activator of macrophages and a systemic inflammatory stimulator, to induce cytokine production. LPS induced the activation of PKD1, which can be detected at 2 h and peaked at 4 h, and returned to baseline at 12 h (Fig. 7C). PKD2 was activated with similar kinetics (Data not shown). The effect of PKD inhibition or depletion on the production of cytokine TGF-β1 was examined upon LPS stimulation. Inhibition of PKD by CRT potently blocked the induction of TGF-β1 by LPS (Fig. 7D). Accordingly, knockdown of PKD1 or PKD2 also partially blocked LPS-induced TGF-β1 production (Fig. 7E). Finally, the effect of PKD inhibition on BMM migration was examined in the presence or absence of CRT. As shown in Fig. 7F, CRT decreased the migratory activity of BMM by 2-fold. Taken together, PKD activity is required for the migration and production of cytokines in macrophages. Our findings imply that PKD may contribute to BLM-induced fibrosis by promoting the recruitment and production of cytokines from macrophages.

**Figure 7.**
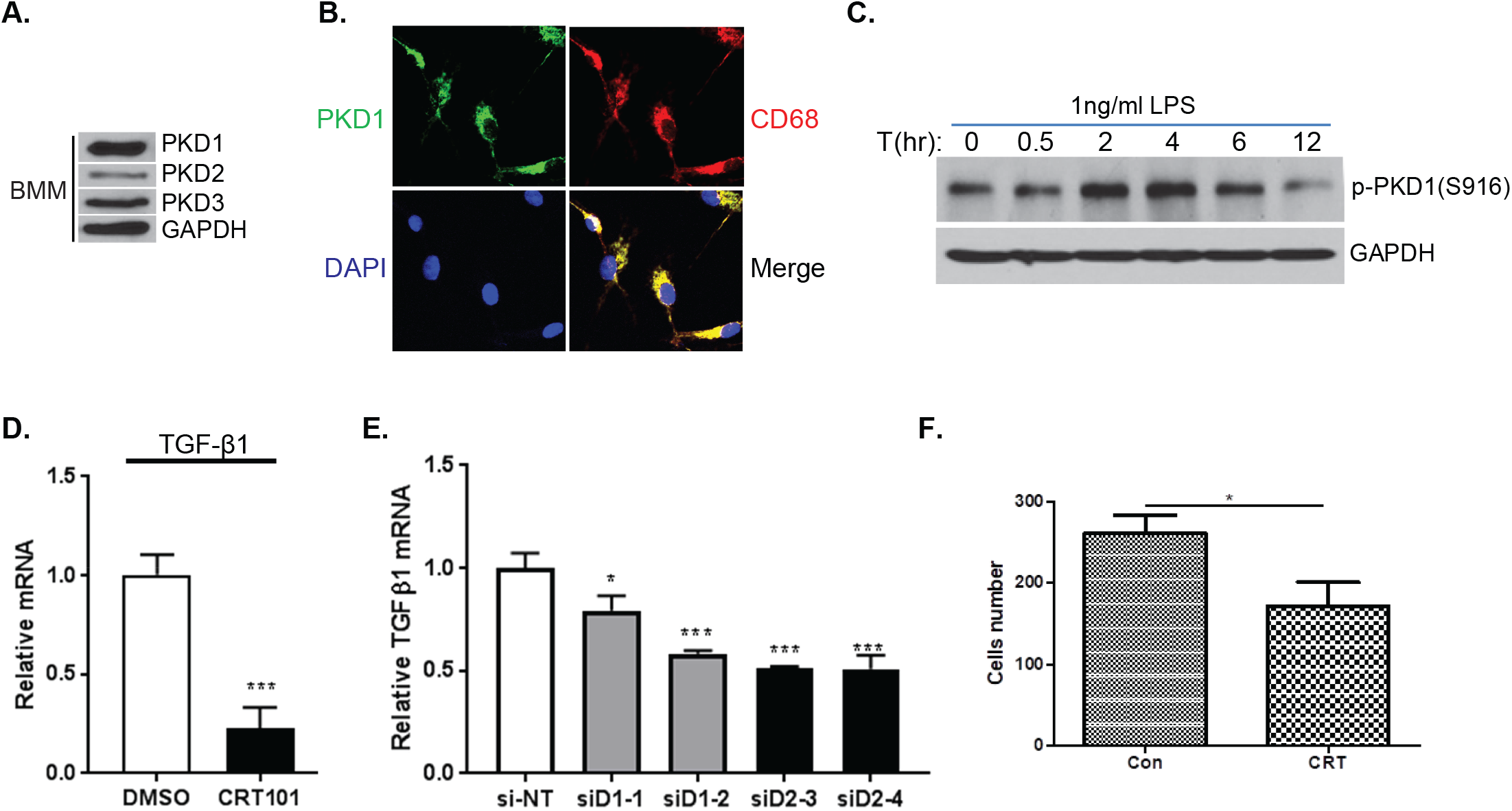
PKD was required for BMM migration and LPS-stimulated cytokine production. (**A**) Expression of PKD isoforms in BMM. Cell lysates of BMM was analyzed by immunoblotting for PKD isoforms. (**B**) BMM was positive for CD68. BMMs were co-stained with PKD1 (green) and CD68 (red), and counterstained with DAPI (blue). Representative images are shown. Original magnification x100. (**C**) LPS activated PKD1. BMM was stimulated with 1 ng/ml LPS for different times. Cells were lysed and subjected to immunoblotting for p-PKD1(S916). (**D**) Effects of CRT on LPS-induced TGF-β1 expression in BMM. BMM were stimulated with 1 ng/ml LPS for 4 h. CRT (2μM) was added in the last 2 h of LPS treatment. Cells were analyzed for TGF-β1 mRNA by real time RT-qPCR. (**E**) Knockdown of PKD1 or PKD2 decreased LPS-induced TGF-β1 expression in BMM. BMM were transfected with PKD1 (siD1-1 and -2) and PKD2 (siD2-1 and -2) siRNAs. Two days after transfection, cells were stimulated with 1 ng/ml LPS for 4 h, and cells were subjected to real time RT-qPCR analysis for TGF-β1 mRNA expression. (**F**) CRT blocked BMM migration.

## Discussion

SSc is a complex multisystem autoimmune disease that has the highest case-specific mortality of the connective tissue illnesses (39). The pathogenesis of the disease is currently understood to involve the interplay of autoimmunity, inflammation, fibrosis, and vasculopathy. Although significant advances have been made in the diagnosis and treatment of SSc, the molecular basis of the disease remains largely elusive and no FDA-approved therapies are available (4, 5). Although PKD has been implicated in fibrosis and inflammation (11, 24, 25, 40), its role in dermal fibrosis involved in SSc has not been investigated. In this study, using a knock-in mouse model of kinase-dead PKD2, we demonstrated a critical role of PKD2 activity in BLM-induced dermal fibrosis.

Dermal fibrosis is characterized by hardening and thickening of dermal layer in skin in part caused by the excessive accumulation of ECM, which contribute to fibrosis development. Activated fibroblasts (myofibroblasts) are known to be the main type of cells responsible for the overproduction of collagens and other ECM components (41). The interactions between fibroblasts and inflammatory cells, and the production of pro-inflammatory cytokines by the inflammatory cells, play a crucial role in the induction of myofibroblastic activity (42). It was reported that TGF-β1 (43) mediates fibroblast proliferation (44), collagen production (45), ECM deposition and myofibroblast differentiation in dermal fibrosis. Enhanced expression of TGF-β1 stimulates collagen synthesis in fibroblasts by activating the smad2/3 signaling pathways (46), resulting in the overabundant accumulation of collagen (47). In addition, at the early stages of SSc, high level of IL-6 was associated with diffuse cutaneous SSc, increased inflammatory markers, more severe skin fibrosis, and worse long-term survival (48).

To determine whether there is a relationship between dermal fibrosis in SSc and PKD2, we obtained homozygous PKD2^SSAA/SSAA^-KI mice where PKD2 activity was eliminated by mutating S707/S711 in the activation loop of PKD2 to define the role of this kinase in BLM-induced dermal fibrosis. Our results suggested that kinase-dead PKD2 could significantly block bleomycin-induced dermal thickening in LS (Fig. 1B, 1C and 1D) and decreased collagen and α-SMA protein expression in mouse model of BLM-induced dermal fibrosis (Fig. 2A, 3A and 4A). Interestingly, as shown in Fig. 2A, 3A and 4A-B, PKD2^SSAA/SSAA^-KI mice also significantly reduced collagen and α-SMA expression under basal state without bleomycin treatment, as compared with PKD2^WT/WT^ mice, suggesting a fundamental role of PKD2 in regulating α-SMA expression. It is known that repeated BLM injections caused infiltration of inflammatory cells, and increased myofibroblastic activity, fibrosis and thickening of the skin, which were observed in our wild-type mice treated with BLM. However, in contrast, PKD2^SSAA/SSAA^-KI mice showed reduced deposition of collagen in the skin, and decreased dermal thickening, and development of fibrosis. We also demonstrated that TGF-β1 and IL-6 mRNA levels were decreased in PKD2^SSAA/SSAA^-KI-BLM mice, relative to PKD2^WT/WT^-BLM mice (Fig. 3B). These results confirm the essential role of the pro-inflammatory and profibrotic properties of PKD2 in BLM-induced dermal fibrosis in mice, implying the PKD may be a potential therapeutic target for skin fibrosis and SSc. Further analysis indicate that PKD2-positive cells were largely myeloid cells that could be recruited to the lesional skin to promote fibrotic process via production of pro-fibrotic cytokines including TGF-β. Future studies on specific PKD isoforms will provide additional insights to the function of this protein kinase family in dermal fibrosis in SSc.

In summary, our results underlie the importance of PKD2 catalytic activity in BLM-induced dermal fibrosis, suggesting that PKD2-positive myeloid cells may contribute to the production of pro-fibrotic cytokines that promote dermal fibrosis induced by BLM. Given our promising data in the mouse model, PKD inhibitors may hold promise for treating SSc.

## List of Abbreviations

SSc: systemic sclerosis
PKD: protein kinase D
PKD2^SSAA^: PKD2^S707A/S711A^
KI: knock-in
BLM: bleomycin
CRT: CRT0066101
TGF-β1: transforming growth factor β1
IL-6: interleukin-6
NBF: neutral buffered formalin
IF: immunofluorescence
BSA: bovine serum albumin
BMM: bone marrow-derived macrophage
H&E: hematoxylin and eosin
IHC: immunohistochemistry
α-SMA: α-smooth muscle actin
LPS: lipopolysaccharide

## Declarations

### Ethics approval and consent to participate

This study was approved by University of Pittsburgh Institutional Animal Care and Use Committee (IACUC).

### Consent for publication

Not applicable.

### Availability of data and materials

All data generated or analyzed during this study are included in this published article [and its supplementary information files].

### Authors’ contributions

QJW and FD conceived and designed the experiments, supervised the experiments, wrote and revised the manuscript.

LC, JZ, YC, AR, WG, JQ, WX, ZX, XL performed experiments and analyzed data.

QJW, FD, LC and WG wrote and revised the manuscript.

BL provided reagents, technical support, general guidance, critical reading and revision of the manuscript.

## Acknowledgements

We are truly thankful for the generous help and guidance we received during the course of this project from Drs. Robert Lafyatis and Robyn Therese Domsic at Department of Medicine, Division of Rheumatology and Clinical Immunology, University of Pittsburgh.

