## Supplemental Fig S1 for "Loss of Protein Kinase D2 Activity Protects Against Bleomycin-induced Dermal Fibrosis in Mice"

### Supplemental Figures

**Figure S1. Generation of the PKD2$^{SSAA/SSAA}$ kinase-dead knock-in (KI) mice.**
(**A**) Depiction of the endogenous mouse Prkd2 allele containing exons 10-18. The black/grey rectangles represent exons and the black arrowheads represent loxP sites. Thick black short lines indicate the positions of the probes were introduced by PCR. The knock-in allele containing the Ser$^{707}$/Ser$^{711}$ mutation in exon 16 is illustrated as a grey rectangle. (**B**) Genotype of PKD2$^{SSAA}$ mutant mice was determined by PCR amplification of genomic DNA. The allele of wild-type mice is a 236bp product, whereas the allele of knock-in mice is a 334bp product. (**C**) PKD2 catalytic activity in different tissues from wild-type and PKD2$^{SSAA/SSAA}$-KI mice was analyzed by western blot using p-PKD2 (ser876) antibody.

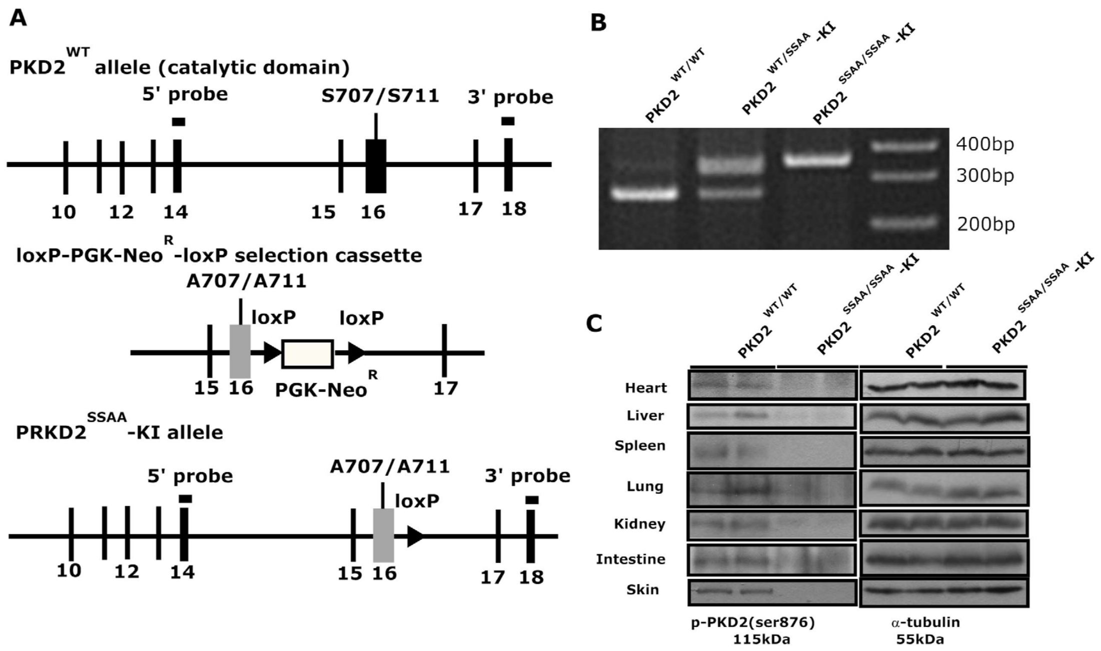
